# On-line Optimization of Hamiltonian Replica Exchange Simulations

**DOI:** 10.1101/228262

**Authors:** Justin L. MacCallum, Mir Ishruna Muniyat, Kari Gaalswyk

**Author notes:** Both authors contributed equally (Mir Ishruna Muniyat, Kari Gaalswyk).

## Abstract

Replica exchange is a widely used sampling strategy in molecular simulation. While a variety of methods exist for optimizing temperature replica exchange, less is known about how to optimize more general Hamiltonian replica exchange simulations. We present an algorithm for the on-line optimization of both temperature and Hamiltonian replica exchange simulations that draws on techniques from the optimization of deep neural networks in machine learning. We optimize a heuristic-based objective function capturing the efficiency of replica exchange. Our approach is general, and has several desirable properties, including: (1) it makes few assumptions about the system of interest; (2) optimization occurs on-line wihout the requirement of pre-simulation; and (3) it readily generalizes to systems where there are multiple control parameters per replica. We explore some general properties of the algorithm on a simple harmonic oscillator system, and demonstrate its effectiveness on a more complex data-guided protein folding simulation.

## Introduction

Replica exchange simulations [1-6] are a widely used sampling technique across a range of disciplines, ranging from molecular simulation [7-12] to Bayesian statistics [13,14]. The relative ease of implementation of replica exchange has lead to its widespread adoption in the molecular simulation community, with implementations available in many major simulation packages [15-20].

Temperature replica exchange, also known as parallel tempering, samples from a series of flattened or tempered distributions, corresponding to a series of increasing temperatures. In biomolecular systems, *N* replicas are simulated across a “ladder” of temperatures, often scaled geometrically from say 300 to 500 K. The simulation proceeds by alternating between normal molecular dynamics or Markov-chain Monte Carlo moves, and exchange moves that attempt to exchange configurations between neighboring replicas. Typically, one is interested in the distribution of configurations of the system at the lowest temperatures, while the higher temperatures allow the system to escape local minima, thus potentially enhancing sampling [6]. One limitation of temperature replica exchange is that efficiency gains are not guaranteed if the barriers to sampling are not dependent on temperature, e.g. entropic barriers [21,22]. Another limitation is that the exchange probability depends on the total potential energy of the system. For large systems, this means that many closely-spaced replicas are needed, which may be computationally prohibitive [8,21,23].

Hamiltonian replica exchange is a more general formulation that uses arbitrary perturbations to the Hamiltonian rather than temperature as the basis for exchange [17, 23-25]. These perturbations can be more targeted, making them potentially more efficient than simply modifying the temperature. One motivating example from work in our lab is integrative structural biology [26], where experimental data—potentially sparse, ambiguous, and unreliable—is used to guide folding [27-29] or binding [30-32] through data-driven restraints. In general, it is possible to combine variations in temperature and multiple different perturbations of the Hamiltonian in a single Hamiltonian replica exchange simulation, where the parameters that determine the Hamiltonian and temperature for each replica are called the *control parameters*.

The efficiency of replica exchange is critically dependent on the values of the control parameters. If the exchange probability between neighboring replicas is too low, it creates a “bottleneck” to replica diffusion that can reduce sampling efficiency. In our view, the difficulty of finding efficient values for the control parameters has been a major impediment to the more widespread adoption of Hamiltonian replica exchange simulations.

The aim of this paper is to develop an online optimization strategy for general Hamiltonian replica exchange. There is extensive literature on the optimization of temperature replica exchange [23,33-36], but far less is known about how to optimize the control parameters for Hamiltonian replica exchange [25,37-39]. While there are several algorithms to optimize control parameters, these often suffer from several drawbacks: (1) they often make assumptions, e.g. that the system has constant heat capacity which may not be true; (2) they often require pre-simulation to gather statistics across the range of control parameters; and (3) most importantly, they are designed for a single control parameter, usually temperature, and do not readily generalize the case of multiple control parameters. Overcoming these limitations is critical to achieving efficient,0 flexible, and general Hamiltonian replica exchange schemes.

In this work, we present an algorithm that avoids these drawbacks. Our approach is to: (1) define an objective function that captures what we mean by efficient replica exchange sampling; (2) compute the derivatives of this objective function with respect to the control parameters so that it can be optimized using gradient-based methods; and (3) perform on-line optimization of the control parameters during the simulation using techniques drawn from machine learning. We examine several properties of the algorithm in a simple harmonic oscillator system, and demonstrate the utility of the algorithm on a data-guided protein folding problem similar to those encountered in integrative structural biology.

## Theory and Methods

### Notation and basic theory of replica exchange

This work considers one-dimensional replica exchange simulations, where a series of replicas are arranged in a “ladder” so that each interior replica has two neighbors. To avoid ambiguity, we use the terminology of *replicas* and *walkers*. Each replica is associated with a Hamiltonian parameterized by a set of control parameters, e.g. temperatures, force constants, etc. A walker is a particular configuration of the system that moves between the replicas through a series of exchange moves, as described below.

The replicas are indexed by *i* = 1… *N*, each with a Hamiltonian, *H_i_*(*x*, **λ**), parameterized by the vector of control parameters, **λ**. We focus on cases where each control parameter affects only a single replica, and where each replica has the same number and type of control parameters, although neither restriction is required. For a system of *N* replicas, each with an associated temperature and force constant, **λ** would consist of *N* temperatures and N force constants, for a total of 2*N* parameters. The control parameters for the top and bottom replica are held fixed.

In general, the Hamiltonians, partition functions, acceptance probabilities, and their averages depend on **λ**. For notational simplicity, we suppress the **λ**-dependence unless it is necessary for clarity. Throughout, we use the reduced Hamiltonian, *h_i_*(*x*) = (*RT_i_*)^−1^*H_i_*(*x*,**λ**), where *T_i_* is the temperature of state *i* (possibly a control parameter), and *R* is the gas constant.

A cycle of replica exchange consists of updating the configuration of each walker by performing a series of molecular dynamics or Markov chain Monte Carlo steps, followed by a series of exchange moves that attempt to swap walkers between randomly selected adjacent replicas. In this work, we consider only exchanges between neighboring replicas, although other strategies are possible [40].

The acceptance probability to swap walker *x_i_* at replica *i* with *x*_*i*+1_ at *i* + 1 is

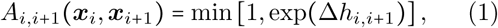

where

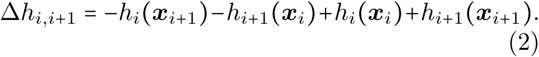

The average acceptance probability between replicas *i* and *i* + 1 is given by an average over the ensembles of both replicas

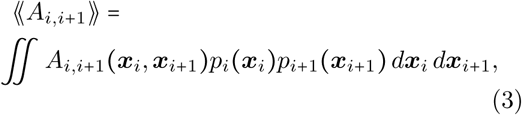

with probabilities 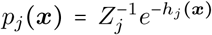 and partition functions *Z_j_* = *∫ e^−h_j(x)_^ d_x_*. We use double angle brackets, 〈〈·〉〉, to indicate averages that depend on neighboring replicas, and single brackets, 〈·〉, to indicate averages that depend on only a single replica. By construction, the average acceptance probabilities are symmetric, so that 〈〈*A*_*i,i*+1_〉〉 = 〈〈*A*_*i*+1,*i*_〉〉.

Throughout, we make the simplifying assumption that the Monte Carlo or molecular dynamics updates between exchanges produce uncorrelated samples from each replica. This implies that various quantities, including the instantaneous acceptance rates, are independent of the history of the system. In practice, this assumption is violated for many systems of interest—usually replica exchange is used because sampling is slow under the conditions of interest, producing highly correlated samples. Thus, our algorithm does not produce truly optimal solutions. Nevertheless, this is a common assumption in the literature [33,36,37,41], and our algorithm appears to be quite useful in practice, as demonstrated below.

### Replica exchange can be optimized using a heuristic

It is surprisingly difficult to quantify exactly what is meant by “optimal sampling” in the context of enhanced sampling algorithms.

Before diving into this question, we note that for replica exchange, there are two broad cases. In the first case, the data obtained at every replica is useful to us, and simulations would be needed at each temperature anyway. Provided that the computational cost of exchange steps is small relative to that of the configuration updates, replica exchange will almost always improve efficiency over independent simulations. In the second case, which is our focus here, we are primarily interested in the results at one replica. In order to gain efficiency, the computational cost associated with the additional replicas must be offset by an increase in sampling efficiency at the replica of interest. Reweighting methods [42,43], which combine the results from multiple replicas to estimate quantities at a single replica of interest, lie between these two extremes.

Returning to the question of optimal sampling, there are two relevant relaxation or correlation times that govern sampling efficiency: (1) how rapidly the sampled distribution decays towards equilibrium from an arbitrary starting state, which is related to the “burn-in” time at the start of a simulation; and (2) how rapidly the sampling algorithm generates statistically independent samples of some function of interest, *f* (*x*), which governs the uncertainty in estimates of 〈*f* (*x*)〉 at equilibrium. Both quantities are directly related to the eigendecomposition of the dynamical operator governing the evolution of the system [44]. Recently, algorithms have been developed for identifying slowly relaxing modes and their associated timescales [44,45], particularly in the context of Markov state models. However, these algorithms are not practical for the optimization of replica exchange simulations because: (1) they require large amounts of simulation data to accurately identify slowly relaxing modes of the system, and (2) it is not straightforward to link changes of the control parameters to perturbations of the eigendecomposition of system dynamics.

Instead, most work aimed at optimizing replica exchange simulations focuses on the heuristic of minimizing the round-trip time [33, 46]—defined as the average time for a walker to transition from the bottom replica to the top, and then back to the bottom again. We adopt the same approach here. Our goal is to sample statistically uncorrelated configurations at the bottom replica. We assume that at the top replica—with the highest temperature, weakest restraints, etc—the walker configuration rapidly decorrelates, effectively “forgetting” its history. Provided the combination of simulation time between exchanges and conditions at the top replica are sufficient to fully decorrelate the walker, each time a walker makes a round-trip, it is guaranteed to produce an independent, uncorrelated sample. The number of round-trips thus sets a lower bound on the number of statistically independent samples produced. Algorithms that optimize round trip time are inherently pessimistic, in that they assume that independent samples of the quantities of interest can only be obtained by completely decorrelating the configuration of the system and, that a round-trip is the only way to achieve said decorrelation.

### The harmonic oscillator is a simple model system

In order to better understand on-line optimization of replica exchange, we initially focus on a classical one-dimensional harmonic oscillator, where the reduced Hamiltonian for replica *i* is given by

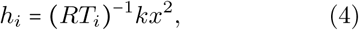

where we fix *k* = 1 kJ mol ^−1^ nm ^−2^, and *T_i_* is a control parameter. For this system, we generate independent, uncorrelated samples between replica exchange steps.

We consider temperature replica exchange with 4 walkers. The first and last temperatures are fixed at *T*_1_ = 10 K and *T*_4_ = 10; 000 K. As a classical harmonic oscillator has constant heat capacity, we expect that an optimal set of temperatures will be geometrically distributed with *T*_2_ = 100 K and *T*_3_ = 1000 K.

### Different objective functions lead to different optimization landscapes

Using the simple harmonic oscillator system described above, we examined four possible objective functions at different combinations of *T*_2_ and *T*_3_ using numerical simulation (Figure 1).

**Figure 1:**
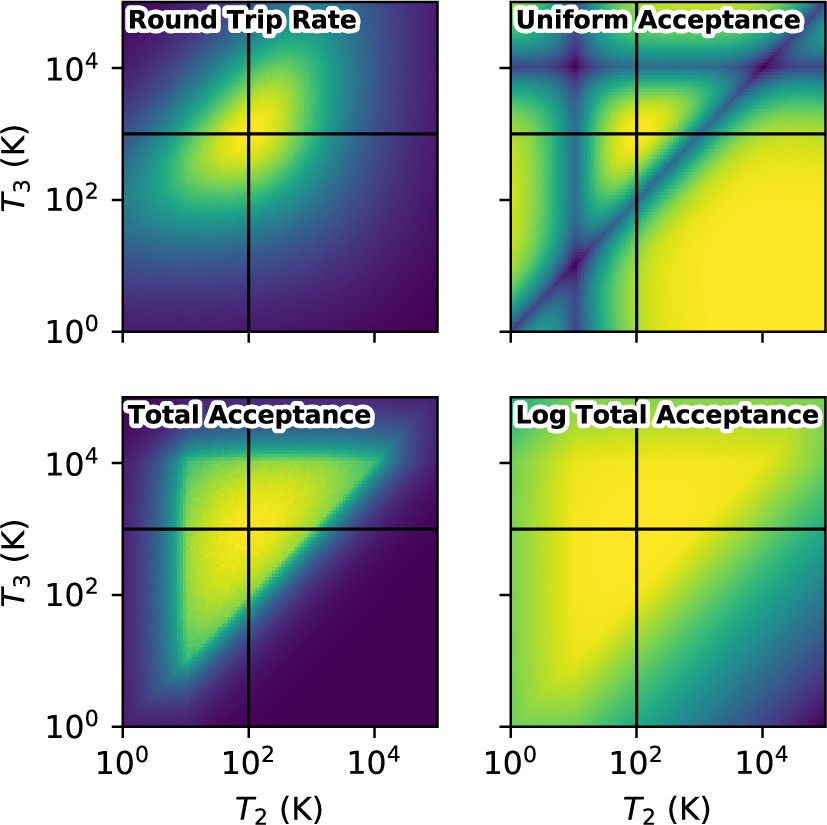
Optimization landscapes for various objective functions as a function of intermediate temperatures *T*_2_ and *T*_3_ with *T*_1_ = 10 K and *T*_4_ = 10; 000 K held fixed. Yellow colors indicate more favorable values of the objective function. The system is a classical harmonic oscillator, as described in the text. The black lines indicate the expected optimal solution.

#### Round trip rate

We first examined the direct use of round-trip rate as an optimization heuristic. There are several approaches to calculating the round-trip rate, but most require either iterative schemes [33] or direct observation of replica flux [46-50]. In either case, it is not immediately clear how to derive an expression for the derivative of the round-trip rate with respect to arbitrary control parameters. Instead, we construct an *N* × *N* transition matrix ***P*** for a random walk through replica space. A walker at replica *i* can jump to *i*−1 with probability ***P***_*i*,*i*−1_ = 0.5 〈〈*A*_*i*,*i*−1_〉〉, jump to *i* + 1 with ***P***_*i*,*i*+1_ = 0.5 〈〈*A*_*i*,*i*+1_〉〉, and remain at *i* with ***P***_*i*,*i*_ = 1 − ***P***_*i*,*i*−1_ − ***P***_*i*,*i*+1_. The mean first passage time matrix ***M*** was obtained by the method of Kemeny and Snell [51]

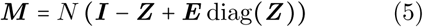

where ***I*** is the *N* × *N* identity matrix, ***E*** is the *N* × *N* matrix of all ones, and ***Z*** is

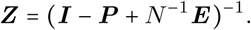

These equations are simplified from those in Kemeny and Snell by the fact that the stationary distribution over replicas is uniform by construction. The objective function is the round-trip rate

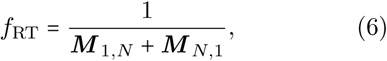

which has an optimum at (*T*_2_ = 100; *T*_3_ = 1000), as expected (Figure 1).

While this objective function directly captures our intent, we observed that it can be unstable when the inverted matrix in the calculation of ***Z*** is singular or nearly so. In particular, for reliable inversion we found that several thousand iterations of replica exchange were required in the calculation of ***P***, which reduces the rate at which adaptation can occur. Although it may be possible to improve the numerical robustness, we did not pursue this objective function further.

#### Uniform acceptance

We next considered an objective based on the heuristic that exchange rates should be uniform

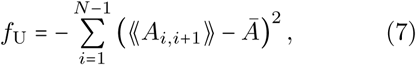

where *A̅* is the average acceptance rate across all pairs of adjacent replicas.

There is a local maximum of this objective function at the expected position (Figure 1). However, there are other local maxima that occur in regions of parameter space where all acceptance probabilities are near zero. For some initial control parameters, *f*_U_ could converge to a good solution, but for other initial control parameters it could converge to a terrible solution. We thus abandoned this objective function.

#### Total acceptance

Following Shenfeld and coworkers [52], we next considered the total acceptance as an objective function

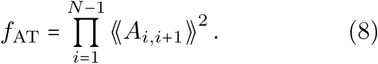

Rather than directly measuring the round-trip rate, this objective function captures the probability of the shortest possible round-trip, where the walker hops from replica 1 to *N* and back in 2*N* – 2 consecutive steps.

This objective function has an optimum in the expected location (Figure 1). However, when the total acceptance is near zero, the gradient of the objective function is minuscule (deep purple region of figure), which is not ideal for the gradient-based optimization strategy we pursue here.

#### Log total acceptance

The poor scaling be-havior of the gradients with the overall magnitude of the objective function can be overcome by using the logarithm of Eq. 8 as the objective function

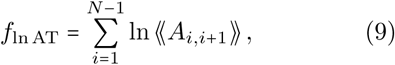

where we have dropped an irrelevant factor of 2.

The optimum of this objective is the same as for Eq. 8, as the logarithm is a concave function. There is a clear gradient in the regions of low total acceptance. However, the objective function is now relatively flat around the optimum. Nevertheless, there is still some gradient in this region, and, as described below, the use of adaptive step sizes and learning rate decay allows for successful optimization.

We use Eq. 9 as the objective function for the remainder of this study.

### Gradient of the objective function

Our approach is to use gradient-based optimization methods developed for machine learning, which require the gradient of the objective function with respect the the control parameters

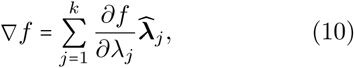

where 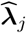 are the basis vectors in the *k*-dimensional space of control parameters.

For the loss function of Eq. 9, the gradient is

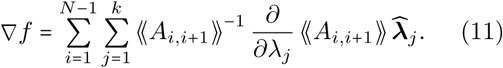

The derivatives are

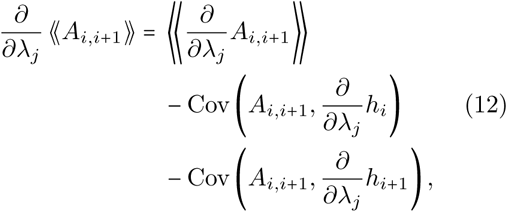

Where

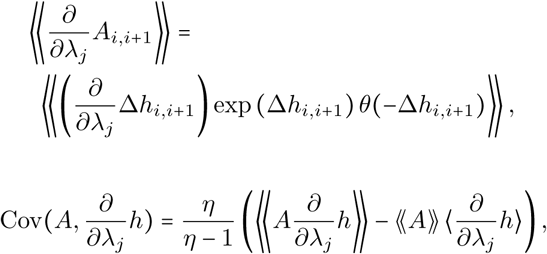

*θ*(·) is the Heaviside step function, and Δ*h*_*i*,*i*+1_ is defined in Eq. 2. The covariances are corrected for bias at small sample sizes, where *η* is the number of replica exchange steps included in the computed averages.

The factor of 〈〈*A*_*i*,*i*+1_〉〉^−1^ in Eq. 11 leads to a singularity when any of the average acceptance probabilities are near zero, which can occur if the initial guess for the control parameters is particularly bad. To mitigate this, we add a small positive constant *∊*_1_ = 10^−9^ to all occurrences of *A*_*i*,*i*+1_ and exp (Δ*h*_*i*,*i*+1_) leading to the following modified expression for the gradient

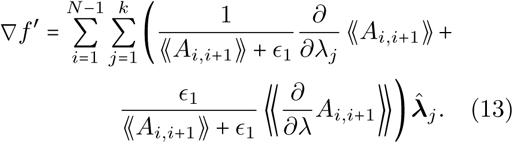

This expression provides an alternative gradient when the acceptance probability is extremely small, making the algorithm more robust.

### The objective function is minimized using stochastic optimization

To optimize Eq. 9, we use the Adam algorithm [53], a stochastic optimization method commonly used to train deep neural networks in machine learning. The parameters are updated according

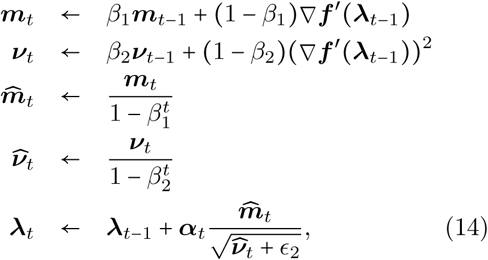

where *m*_0_ = *v*_0_ = 0, is updated after each adaptation step by *t*←*t*+1, *α_t_* is the learning rate (discussed below), **λ**_0_ is the initial parameter guess. We set the constants *∊*_2_ = 10^−9^ and *β*_1_ = *β*_2_ = 0.9, as we found that the parameters suggested for machine learning applications, (*β*_1_ = 0.9, *β*_2_ = 0.999), did not adjust quickly enough to the noise and sudden changes in gradient encountered in our application.

### Schedules of learning rate and averaging time allow for efficient optimization

To allow the system to settle into an optimum in the presence of noisy estimates of the gradient, the learning rates are reduced throughout the run by

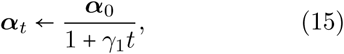

where *γ*_1_ is the learning rate decay parameter. The initial learning rate vector, *α*_0_, is system-dependent. In this work, we set the initial learning rates for all parameters of a given type to be the same, e.g. the temperature parameters for all replicas share the same learning rate. The learning rates for the top and bottom replicas are set to zero.

To compute the gradient, the ensemble averages in Eqs. 11 and 12 are evaluated over a number of replica exchange steps, *η*, called the averaging time. When *η* is small, the gradients are noisy, which prevents the parameters from settling into an optimum. Conversely, when *η* is large, the gradients are less noisy, but this comes at the cost of fewer adaptation steps per unit of simulation time. In this work, we increase the averaging time throughout the simulation by

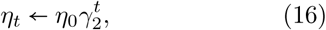

where *η*_0_ is the initial averaging time and *γ*_2_ is the averaging time growth.

After a change in parameters, the system will temporarily be out of equilibrium, causing the average gradient to deviate from the correct value. To mitigate this, we adapt the parameters every 2*η_t_* steps, where the first *η_t_* steps are discarded and the remainder are used to compute the gradient.

Together, the parameters for the optimization algorithm, ***λ***_0_, ***α***_0_, *η*_0_, *β*_1_, *β*_2_, *γ*_1_, and *γ*_2_ are referred to as *hyper-parameters*.

### Several criteria can be used to terminate optimization

There are several possible criteria to decide when to stop optimization. (1) Optimization may not be terminated at all, with constant updates to parameters throughout the simulation. This may be acceptable in some cases due to the decreasing frequency and magnitude of parameter updates owing to the averaging time growth and learning rate decay. However, for critical applications, even small changes to the parameters and associated non-stationarity of the distributions may be unacceptable. (2) Optimization may be stopped after a fixed number of steps. (3) Optimization may be stopped after the change in parameters between successive updates drops below a particular threshold. (4) Optimization may be stopped once all average exchange rates lie between 7% and 82%, as it is known that under these conditions round-trip rates for optimal and near-optimal solutions can differ by a factor of 2 at most [33]. There are diminishing returns in further optimization compared to increasing the amount of sampling with fixed, if slightly sub-optimal, parameters. In this work, we use option 1, as we are interested in the behavior of the algorithm, rather than the actual results of the simulation. For general use, we recommend either option 3 or 4.

### Data-guided protein folding is a more complex model system

For a more complex model system, we examined the data-guided folding of Protein G. This test is designed to mimic the challenges encountered in integrative structural biology applications [26], where the task is to use data that may be sparse, ambiguous, or unreliable [27-29, 54, 55] to guide protein folding.

We focus on the data-guided folding of Protein G using native-centric backbone dihedral and C_*α*_–C_*α*_ restraints. This is not meant to be particularly realistic—all of the restraints are derived from a previously published NMR structure [56], they are accurate, and they are not particularly sparse (Figure 2). Nevertheless, we have found that, even for such simple systems, it is difficult to obtain a good set of parameters for efficient Hamiltonian exchange.

**Figure 2:**
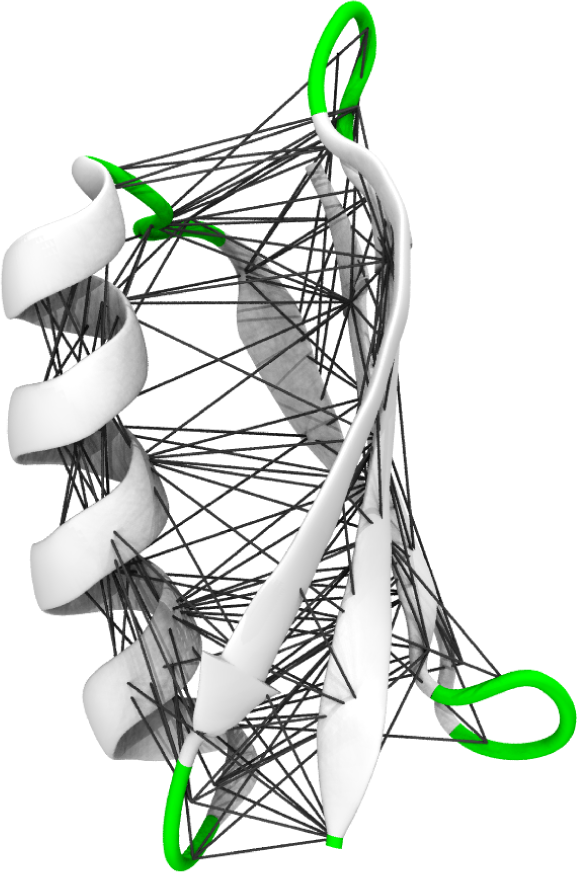
Restraints used during data-guided protein folding simulations. The secondary structures in the white regions were restrained to the correct backbone *ϕ*/*ψ* angles, whereas green regions were unrestrained. The black lines indicate C_*α*_–C_*α*_ distances that were restrained to the experimentally observed values. No restraints were applied to any of the side chain atoms.

The system was modeled using the ff14SB [57] force field with the OBC implicit solvent model [58] and a grid-based correction to the backbone *ϕ*/*ψ* potential to better reproduce secondary structure propensities [59]. Simulations were carried out using the OpenMM library [60, 61], version 7.1. Backbone dihedral angles within secondary structures were weakly restrained to their native values with a force constant of 20 kJ mol^−1^ rad^−1^. The strength of these restraints was held constant across replicas. Restraints were added for all pairs of C_*α*_ with native distances below 10 Å with a force constant that varied across replicas, *k_i_* ∊ [*e*^−10^, 250] kJ mol^−1^ nm^−2^. The temperature was also varied across replicas, *T_i_* ∊ [300,450] K. We used *T_i_* and ln *k_i_* as control parameters.

We examined two different sets of initial parameters, which we refer to as the *simultaneous* and *separate* cases. All simulations used 16 replicas. For the simultaneous case, *T_i_* and ln *k_i_* are varied together linearly across the entire range of replicas. For the separate case, *T_i_* varied linearly from 300 to 450 K across replicas 1-8, while ln *k_i_* varied linearly from −10 to ln 250 kJ mol^−1^ nm^−2^ across replicas 9-16. In all cases, we used an initial guess of hyper-parameters motivated by physical intuition, with 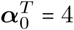, 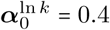, *η*_0_ = 16, *γ*_1_ = 1/100, and *γ*_2_ = 2^1/100^.

## Results

### Learning rate and averaging time affect optimization

Figure 3 shows the influence of *α* and *η* on optimization for the simple harmonic oscillator system. In this experiment, the learning rate and averaging time are held constant in each simulation with *γ*_1_ = 0 and *γ*_2_ = 1.

**Figure 3:**
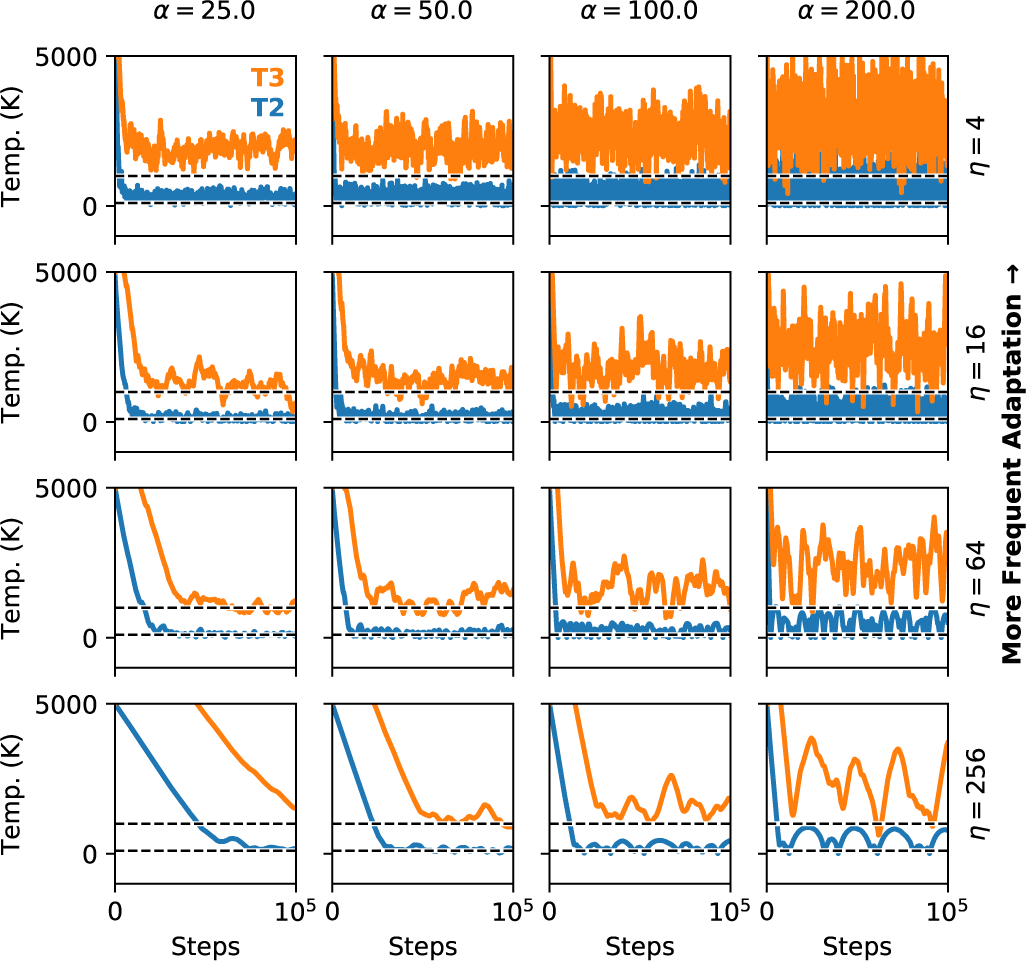
Effect of learning rate (*α*) and averaging time (*η*) on the optimization of temperature replica exchange for a simple harmonic oscillator. *T*_2_ and *T*_3_ are initially set to 5000K, while *T*_1_ = 10 and *T*_4_ = 10,000 K are fixed. The learning rate and averaging time are held constant with *γ*_1_ = 0 and *γ*_2_ = 1. The dashed lines indicate the expected solution.

The initial transient decay from the initial parameters is more rapid at higher learning rates. However, there is also more noise and the longterm fluctuations are larger. Low learning rates have more stable long-term behavior, but the initial transient response is much slower.

Similar observations can be made for the averaging time. When the averaging time is short, the results are noisy and the system does not settle into a well-defined optimum. The presence of noise in the gradients is akin to temperature in simulated annealing, and it has been suggested that adding noise to the gradients can be helpful for escaping local minima in machine learning [62,63]. When the averaging time is long, the long-term fluctuations are smaller, but it takes substantially longer to reach the optimal value.

It is evident that results systematically deviate from the optimum when *η* is small (top row of Figure 3; bottom row of Figure 4). This occurs because the estimator of the gradients in Eq. 11 is biased for small *η*, but appears to be asymptotically unbiased.

**Figure 4:**
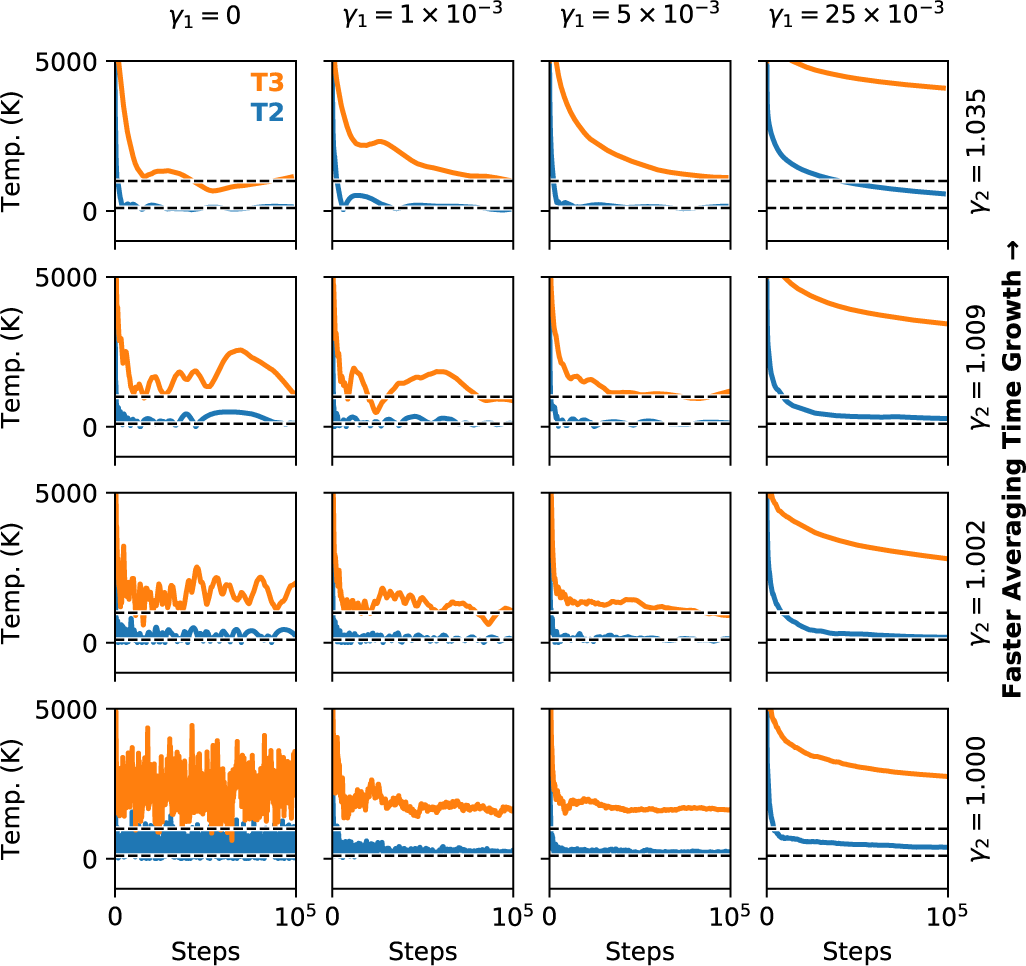
Effect of learning rate decay (*γ*_1_) and averaging time grown (*γ*_2_) on the optimization of temperature replica exchange for a simple harmonic oscillator. *T*_2_ and *T*_3_ are initially set to 5000K, while *T*_1_ = 10 and *T*_4_ = 10,000 K are fixed. The initial learning rate and averaging time are set to *α*_0_ = 100 and *η*_0_ = 4, respectively. The dashed lines indicate the expected solution.

Figure 4 shows the influence of the learning rate decay (*γ*_1_) and averaging time growth (*γ*_2_) on optimization for the simple harmonic oscillator system. Extreme values of either parameter result in large fluctuations or slow decay towards the optimum. However, a variety of combinations result in reasonable behaviour.

Overall, the ideal hyper-parameters are system specific. In this work, we set these hyperparameters guiding by biophysical intuition. In our experience, it is straightforward to find values that work sufficiently well. It is also possible to use a grid search or randomized hyper-parameter optimization. We expect that hyper-parameters should generally be transferable between similar systems.

### On-line adaptation of replica exchange can optimize data-guided protein folding simulations

Our algorithm was able to successfully optimize replica exchange data-guided protein folding simulations. For the *separate* initial parameters, there were large regions of the replica exchange ladder where the initial exchange rate was poor (Figure 5, upper-left). During the course of the simulation, exchange rates became more uniform and overall diffusivity of replicas was improved dramatically (Figure 5, upper right). We observed similar behavior for the *simultaneous* initial parameters (not shown). Although the *separate* initial parameters varied the temperature only over replicas 1–8, the optimized parameters vary the temperature over the full 16 replicas with the temperatures “spreading out” over the course of the optimization (Figure 6). Similarly, although the changes to ln *k* were initially confined to the top 8 replicas, after optimization, the changes are distributed more evenly across replicas.

**Figure 5:**
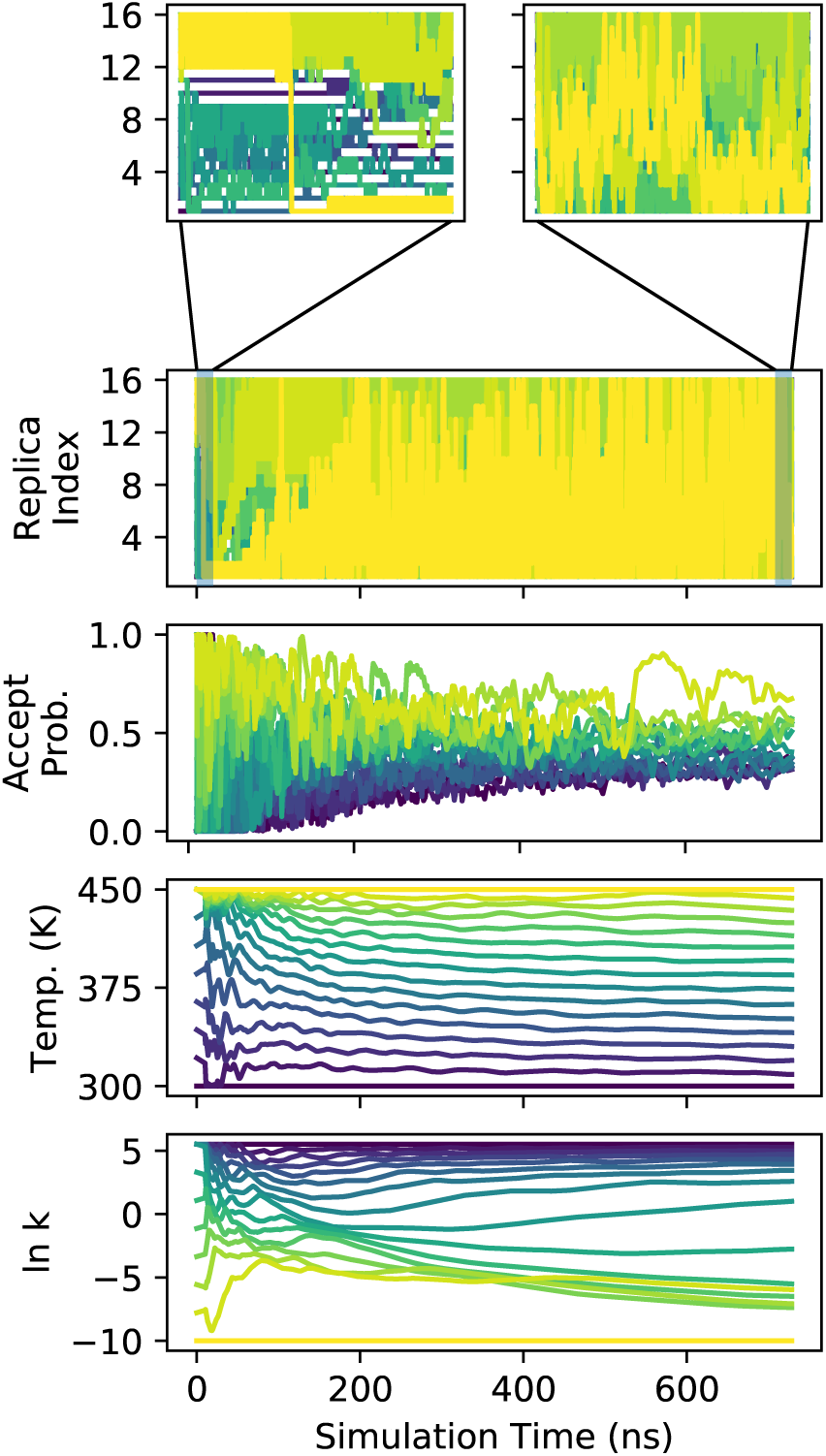
Visualization of replica exchange adaptation for data-guided protein folding starting from the *separate* initial conditions. Colors indicate replica index, from 1 (purple) to 16 (yellow). The top panels show detailed time traces over the first and last 10 ns of the simulation.

**Figure 6:**
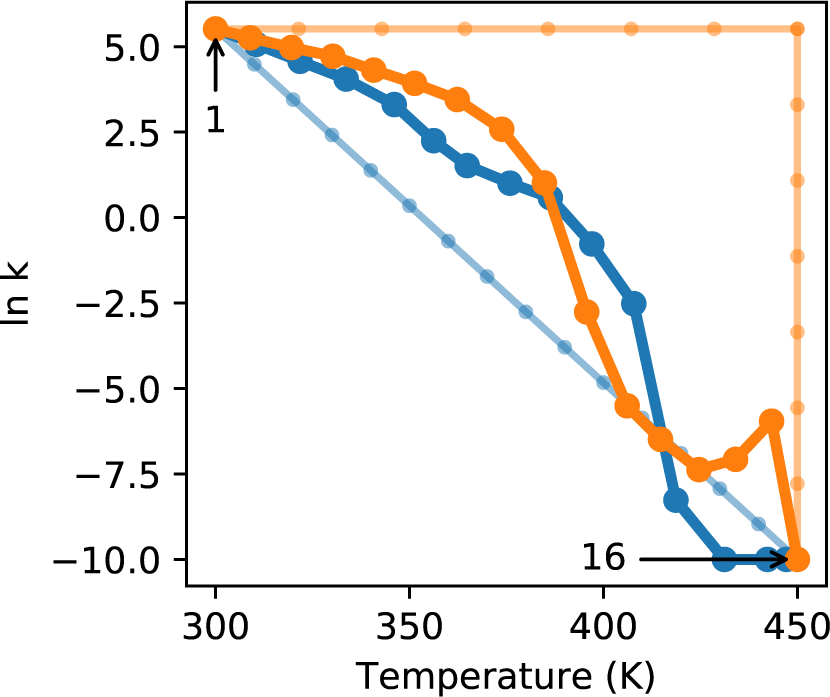
Visualization of initial (faded colors) and final (dark colors) adapted parameters for data-guided protein folding simulations for the *separate* (orange) and *simultaneous* (blue) initial parameters. Numbers indicate the first (1) and last (16) replicas.

The parameters, particularly ln *k*, are still slowly drifting after 700 ns (Figure 5), which may indicate that the optimal solution has not yet been reached. These results are from a single guess for the hyper-parameters of the algorithm, guided primarily by our physical intuition, and it is possible that a more extensive hyper-parameter search could lead to more rapid optimization. However, as noted previously, acceptance rates can only improve by at most a factor of 2 once all acceptance probabilities fall between 7& and 82% [33]. In practice, the most efficient strategy would be to terminate optimization once this occurs, which in this case would be after ~ 200 ns, resulting in ~200 ns of optimization and 500 ns of production simulation.

Figure 6 compares the optimized solutions starting from either the *separate* or *simultaneous* initial parameters. Although the initial parameters differ dramatically, the final optimized solutions are quite similar. In both cases, the optimized temperature change occurs over the full range of replicas. The behavior of ln *k* is slightly more complex, with two different regimes evident. For the *separate* initial parameters, there is a “spike” in ln *k* over the last few replicas, however this occurs when the force constant is very small, ln *k* = −5 ⟹ *k* = 7 × 10^−3^, and thus likely has little influence on the simulation.

As would be expected from the native-centric restraints used to guide folding, the structures generated by data-guided folding are highly accurate (Figure 7). For both the *separate* and *simultaneous* initial conditions, the backbone root mean square deviation is below 1 Å for nearly all structures sampled after the first ~ 200 ns (not shown). More impressive is the near-perfect superposition of side chain conformations, which were completely unrestrained. Through the simulation, walkers are unfolded by round-trips to high temperatures. The side chains are able to correctly repack during the data-guided refolding of the backbone. This is a demonstration of the power of physics-based integrative structural biology, where the use of data to guide certain features of the structure leads to accurate predictions, even for features that are unrestrained by data.

**Figure 7:**
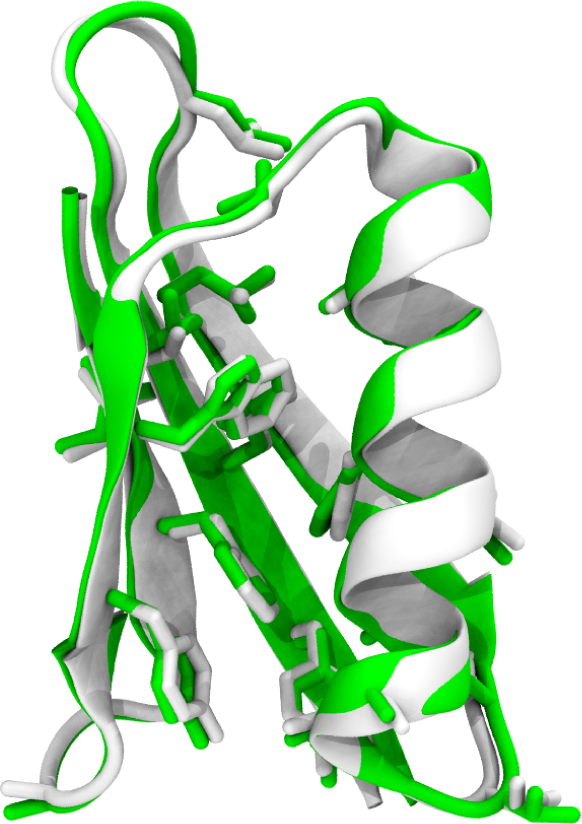
Final frame of data-guided protein folding simulation (green) compared to the reference structure (white; PDB identifier 3gb1). The core side chains are in near-perfect superposition.

## Discussion and Conclusions

We have presented a general framework for the online optimization of Hamiltonian replica exchange simulations. Our algorithm is able to successfully optimize replica exchange for a restrained protein system, reminiscent of those encountered in integrative structural biology.

Our algorithm has three unique properties compared to existing approaches [23,25,33-39]. First, it is an on-line algorithm. Given a suitable criterion to terminate optimization, it is possible to both optimize replica exchange and collect equilibrium statistics from a single simulation. Second, our algorithm straightforwardly handles arbitrary modifications of the Hamiltonian, extending beyond temperature replica exchange. This should help to enable new Hamiltonian replica exchange sampling schemes, which have great promise, but are often difficult to tune in practice. Third, our algorithm readily handles multiple control parameters per replica. This is a major advantage, as it allows for multiple perturbations of the Hamiltonian to be combined. In the case of integrative structural biology, we might combine temperature replica exchange with multiple types of data-driven restraints. The optimal set of control parameters is likely system- and data-dependent and difficult to predict, making such simulations difficult without a strategy for optimization.

Our algorithm does have some potential shortcomings. First, optimization requires the specification of several hyper-parameters, and inappropriate choices of these parameters may lead to poor optimization. Nevertheless, we have found it straightforward to find reasonably well-performing hyper-parameters through simple trial and error.

A second major shortcoming is that our algorithm assumes that the sampling between replica exchange steps leads to uncorrelated configurations. In practice, this is almost never the case, as replica exchange is primarily used when relaxation is slow and sampling is difficult—leading to highly correlated configurations. This means that, although our algorithm is able to generate reasonable control parameter values starting from an arbitrary initial guess, it may be possible to identify parameters that produce even better sampling. Indeed, there are several papers— based on variations of a single algorithm—that demonstrate this [34,35,46-50,64]. However, these methods face several challenges of their own. (1) These algorithms inherently depend on global properties of the system, where each walker must visit each replica multiple times before optimization can proceed. In cases where mixing is slow, this can make reliable optimization prohibitively expensive. (2) Anecdotally, we have found that these algorithms can be difficult to converge, as the optimal solution depends on a delicate balance between overall acceptance rate and more closely spacing replicas in regions of parameter space where relaxation is slow. (3) These approaches do not address the important case of multiple control parameters per replica, which is potentially a major limitation for Hamiltonian replica exchange. We believe that the ideas presented in our work can be extended to account for slow relaxation, and work along these lines is underway in our laboratory.

Hamiltonian replica exchange is a powerful sampling strategy that has yet to reach its full potential, in part due to the difficulty of finding appropriate paths through the space of control parameters. The algorithm presented here provides a robust way to optimize these control parameters, which leads to more efficient simulations. Our approach should help enable the development of new Hamiltonian replica exchange schemes.

## Acknowledgements

This work is supported by funding from the Natural Sciences and Engineering Research Council of Canada and the Canada Foundation for Innovation, and by computational resources from Compute Canada. JLM is a Tier 2 Canada Research Chair. JLM thanks David Sivak and Steven Large for fruitful discussions.

